# The effect of APOBEC3B deaminase on double-stranded DNA

**DOI:** 10.1101/750877

**Authors:** Joseph H. Chapman, Michael F. Custance, Gianna M. Tricola, Birong Shen, Anthony V. Furano

## Abstract

Mutations mediated by the APOBEC3 (A3) family of single-strand specific cytosine (C) deaminases can accumulate as strand-coordinated clusters and isolated lesions during development and in many cancers. A3B is a member of this family with a strong preference for C in a T<u>C</u> context that catalyzes hydrolysis of the primary amine of unpaired C to generate uracil (U). Although single-stranded DNA is the preferred A3B substrate, we report here that compromised hydrogen bonding of C in double-stranded DNA renders it susceptible to A3B deamination *in vitro*. We show that A3B can deaminate the C of T<u>C</u> in a dsDNA context *in vitro* when C is opposite an O^6^-methylguanine or an abasic site. These novel substrates could explain the origin of previously described genomic A3-mediated isolated mutations. We also show that elevated expression of A3B can enhance double-strand breaks induced by the guanine methylating agent, MNNG, in mammalian cells, which is independent of A3B deaminase activity.

## Introduction

Humans encode seven APOBEC3 (A3) single-strand specific cytosine deaminases, six of which (A3A, A3B, A3C, A3D, A3F, A3H) prefer C in a T<u>C</u> context,, while the 7^th^, A3G, prefers CC^1^. Numerous cancers (including head, neck, bladder, lung, cervical, ovarian, and breast) can accumulate A3-mediated mutations - defined as mutations of C in a T<u>C</u> context to similar amounts of transitions (C-to-T) and transversions (C-to-A and C-to-G)^2–6^. A3-mediated mutations can occur as isolated lesions or in strand-coordinated clusters^6–8^ and A3B, in particular, has been implicated in the mutagenesis of several human cancers^2–4^. Such mutations can also occur in normal cells and account for ~20% of heritable mutations^9^.

Strand-coordinated clusters of A3B mutations can result from single-stranded DNA (ssDNA) generated during transcription, replication, recombination, and repair, but the origin of isolated A3B mutations remains unclear^2–4,9–11^. We hypothesized that such isolated mutations could arise in double-stranded DNA (dsDNA) where a C in a T<u>C</u> context is opposite an O^6^-methylguanine (O^6^meG) or an abasic (AP) site, DNA lesions that are not that uncommon.

AP sites, which are potentially mutagenic^12^, are the most common genetic lesions^13,14^ and can result from spontaneous depurination, reactive oxygen species (ROS), or as a consequence of the first step of base excision repair (BER) via glycosylase-mediated removal of abnormal or mismatched bases^13,15–18^. U/G and T/G mispairs, due respectively to hydrolytic deamination of C or 5-methyl-C, are major BER substrates and contribute to upwards of 10,000 AP sites per mammalian cell produced daily, though this number can vary widely between tissues^13,14,18,19^.

O^6^meG is highly mutagenic and carcinogenic^20^ and is produced by alkylating agents such as MNNG^21^, chemotherapeutic agents^22^, or naturally occurring nitrosamines^23^. Nitrosamines are found in processed food and are generated *in vivo* by nitric oxides^24^. About 1.1-6.7 adducts per 10^7^ guanine residues occur in the liver and 0.7-4.6 adducts per 10^8^ guanine residues in leukocytes^13,25^. O^6^meG can be repaired by O^6^-methylguanine-DNA methyltransferase (MGMT) via transfer of the O^6^-methyl to the cysteine (Cys145) in the MGMT active site, restoring the G and inactivating MGMT^26–30^. Persistence of O^6^meG can lead to double-strand breaks (DSBs)^31^.

Here we report that compromised hydrogen bonding of C in dsDNA, specifically C opposite O^6^meG or an AP site, renders it susceptible to A3B deamination *in vitro.* We also found that A3B can enhance DNA damage caused by O^6^meG in mammalian cells, although this effect is independent of its deaminase activity.

## Results

### C in a T<u>C</u>-context opposite O^6^meG or an AP site in dsDNA is susceptible to A3B-catalyzed deamination

We hypothesized that compromised hydrogen bonding of the primary amine of C paired to O^6^meG could make it susceptible to A3B-catalyzed deamination (Fig. **1a**). To test this, we used an *in vitro* A3B deamination assay (Fig. **1b**) by treating an annealed 39 bp oligonucleotide containing an O^6^meG/C base pair (Supplementary Table **S1**) with an extract of HEK293T cells that had been transfected with an A3B-HA expression plasmid (Supplementary Fig. **S1**, insert). The DNA strand containing the putative A3B-susceptible C is 5’ labeled with fluorescein. A3B deamination of C produces U, a substrate for uracil-DNA glycosylase (UDG), which generates an AP site that is labile to NaOH hydrolysis. The ensuing cleaved product is separated from the intact substrate by denaturing urea polyacrylamide gel electrophoresis (Urea PAGE) (Fig. **1b**). Electrophoresis on a native polyacrylamide gel showed that >98% of the annealed O^6^meG/C oligonucleotides in both T<u>C</u>T and A<u>C</u>T contexts were double-stranded (Fig. **1c**). The extent of deamination on the TCT-containing dsDNA substrates (except T<u>C</u>T/AGA) was a function of the amount of HEK293T extract, reaching a maximum at ~25μg (Supplementary Fig. **S1**), which was the amount that we routinely used. The *in vitro* deamination assay showed that O^6^meG/C in a T<u>C</u>T but not A<u>C</u>T context can be deaminated by HEK293T extract containing A3B-HA (Fig. **1d**).

**Fig. 1:**
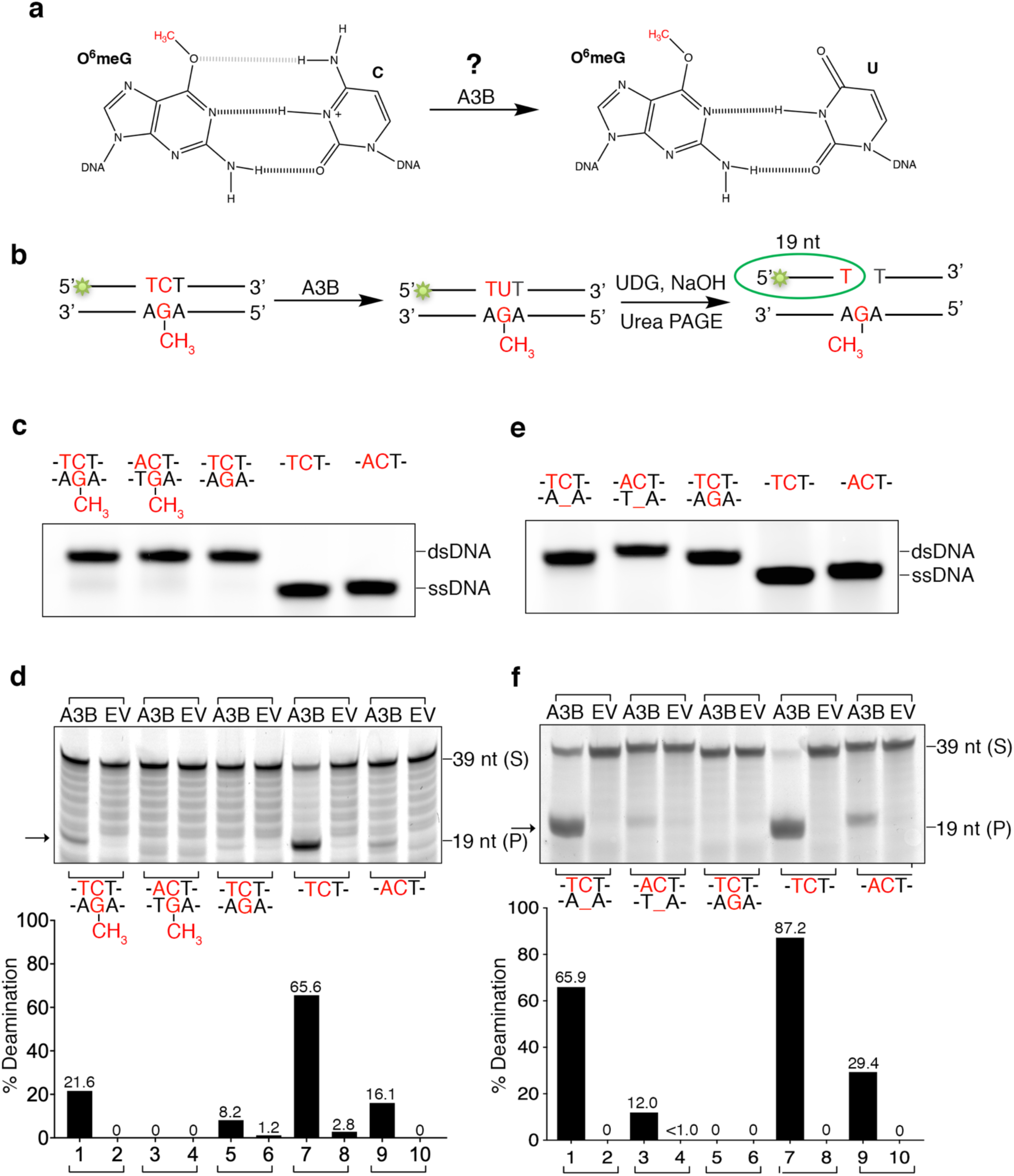
Cytosines (in a TC context) opposite O^6^meG or AP sites are A3B substrates in vitro. (**a**) Hypothesis of O^6^meG in double-stranded DNA as an A3B target. The weakened O^6^-G/N^4^-C hydrogen bond (gray) allows A3B to deaminate C, resulting in an O^6^meG/U base pair. (**b**) Diagram of the *in vitro* deamination assay. (**c,e**) O^6^meG/C (**c**) or AP/C (**e**) 39 bp oligonucleotides run on a native 12% polyacrylamide gel to verify DNA annealing. Single- and double-stranded oligonucleotide serve as controls. (**d,f**) *In vitro* deamination of overexpressed A3B-containing HEK293T whole cell extract on O^6^meG/C (**d**) or AP/C (**f**). TC-containing ssDNA serves as a positive control, and AC-containing ssDNA and TC-containing dsDNA serve as negative controls. Bar graphs (lower panels) show the amount of cleaved product relative to the amount of starting substrate. A3B, HEK293T whole cell extract containing overexpressed A3B-HA; EV, HEK293T whole cell extract that had been transfected with an empty vector. S, substrate; P, cleaved product.

Likewise, an AP/C base pair would expose the C-amine group to a greater extent and could also be an A3B substrate. Both AP/C oligonucleotides are >99.9% double-stranded as shown by native PAGE (Fig. **1e**), and AP/C in a T<u>C</u>T but not A<u>C</u>T context can be deaminated by A3B-HA-containing HEK293T cell extract (Fig. **1f**). Thus, T<u>C</u> opposite O^6^meG and AP sites in dsDNA are A3B substrates *in vitro*.

### Hairpin oligonucleotides containing O^6^meG are vulnerable to A3B-catalyzed deamination

Although our results show that the annealed O^6^meG/C double-stranded oligonucleotide is an A3B substrate (Fig. **1d**), we observed a trace amount of unannealed T<u>C</u>T strand (~1.7%, Fig. **1c**, lane 1). This could not account for the ~21.6% deamination product (Fig. **1d**, lane 1). Nonetheless, we examined deamination of C opposite O^6^meG in an assuredly double-stranded context by imbedding an O^6^meG/C or O^6^meG/U (positive control) in a 182 nt hairpin oligonucleotide (Fig. **2a-b**). Similar to the annealed double-stranded oligonucleotide, the O^6^meG/C hairpin oligonucleotide was deaminated by A3B-HA-containing HEK293T cell extract (Fig. **2c**).

**Fig. 2:**
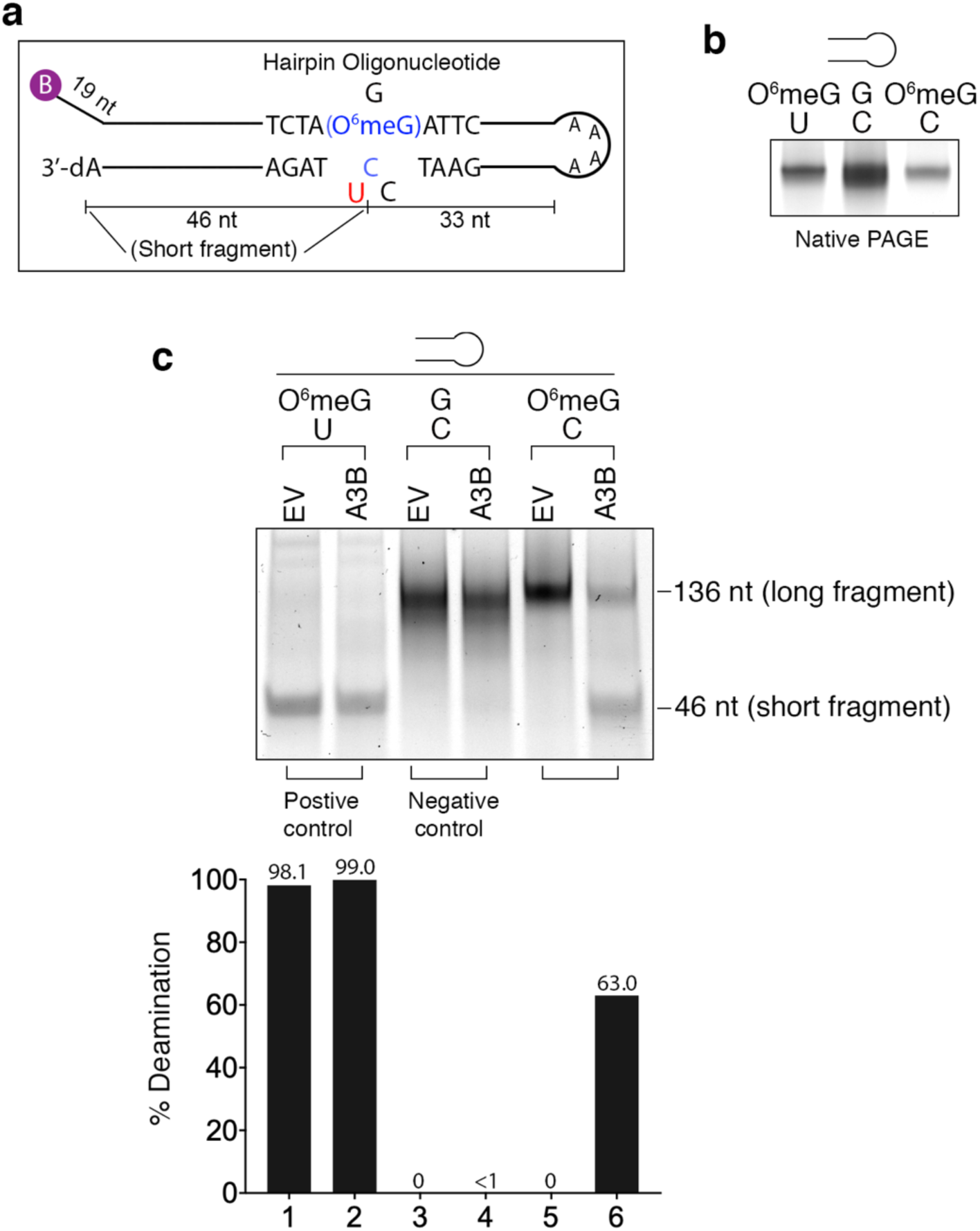
Cytosine (in a TC context) opposite O^6^meG in a hairpin is an A3B substrate in vitro. (**a**) Diagram of the 3 hairpins used in these experiments - O^6^meG/U, G/C, and O^6^meG/C. (**b**) Hairpin oligonucleotides run on a native 12% polyacrylamide gel and stained with GelRed. (**c**) *In vitro* deamination with overexpressed A3B-containing HEK293T whole cell extract on O^6^meG/U, G/C, or O^6^meG/C-containing hairpin oligonucleotides. 12% urea polyacrylamide gel was stained with GelRed to visualize DNA. Bar graph (lower panel) shows the amount of cleaved product relative to the amount of starting substrate. A3B, HEK293T whole cell extract containing overexpressed A3B-HA; EV, HEK293T whole cell extract that had been transfected with an empty vector.

### A3B is required and the carboxy terminal domain of A3B (A3B-CTD) is sufficient to deaminate lesion-containing dsDNA in vitro

The requirement for a T<u>C</u> context strongly suggests that deamination of C is due to A3B activity in HEK293T extracts. We verified this by showing that extracts, which contain A3B-HA mutated at positions in or near the A3B active site (based on previous work by Shi et al.^32^, Fig **3**, upper panel, Supplementary Table **S2**) eliminated or significantly reduced (A254G or S282A) A3B deaminase activity on the T<u>C</u>T single strand, AP/C, and O^6^meG/C oligonucleotides (Fig. **3**, lower panels).

**Fig. 3:**
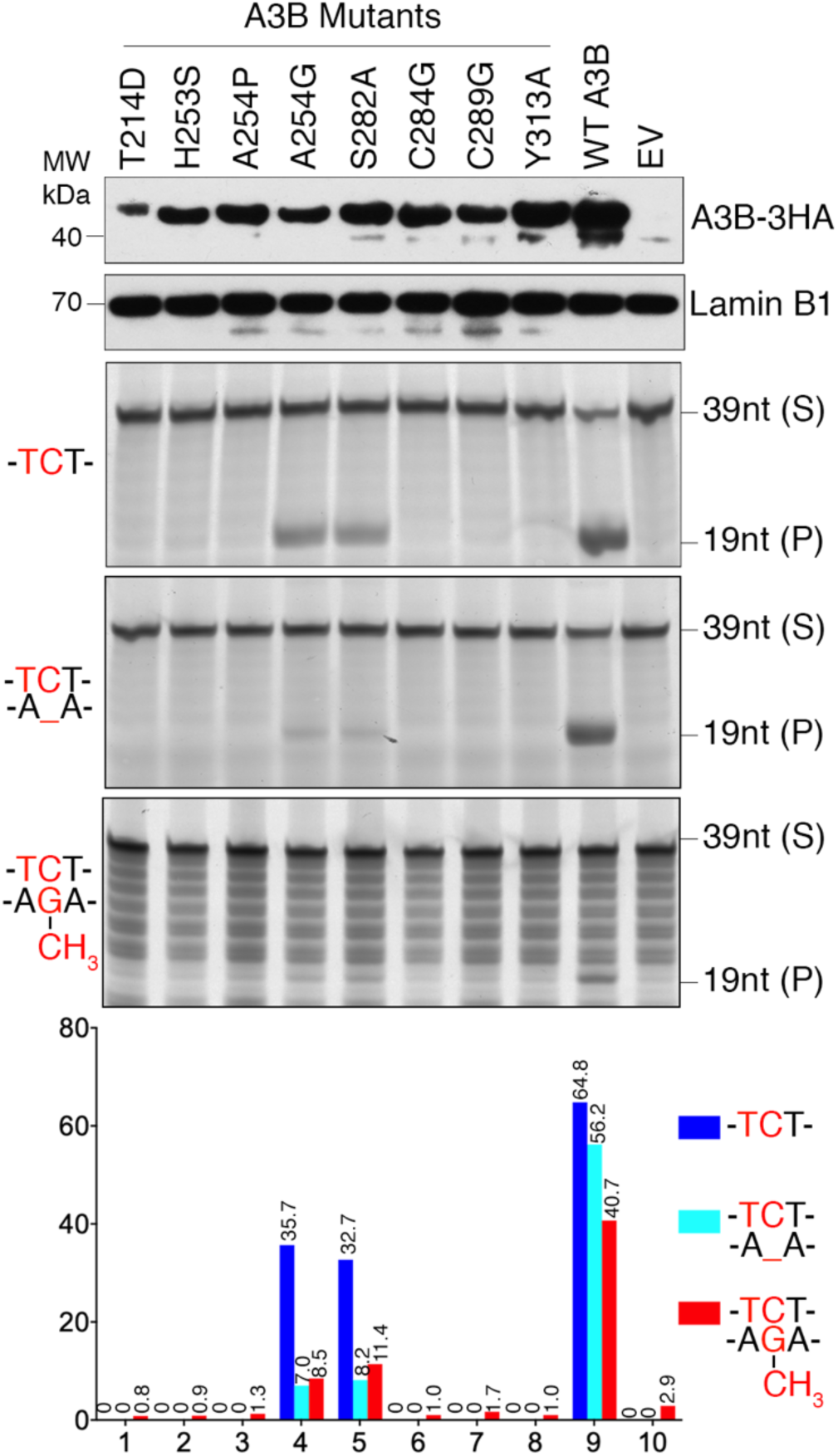
Deaminase activity is reduced using A3B mutant proteins. Western blots of HEK293T nuclear extracts from cells transfected with expression vectors containing various mutant A3B-HA proteins probed with anti-HA antibody (top panel). LaminB1 serves as a loading control (second panel). *In vitro* deamination with A3B mutant protein-containing HEK293T nuclear extract on O^6^meG/C, AP/C, or TC-containing ssDNA (bottom 3 panels). Bar graphs show the amount of cleaved product relative to the amount of starting substrate. WT, wild type; S, substrate; P, cleaved product.

To corroborate the above conclusion, we carried out *in vitro* deamination assays with highly purified A3B-CTD (Fig. **4a**), which exhibits T<u>C</u>-specific deaminase activity^33–35^. We showed that 10 μM of purified A3B-CTD (Fig. **4a**) is sufficient to deaminate ~70% of single-stranded T<u>C</u>T oligonucleotides (Fig. **4b**). A3B-CTD could deaminate C opposite either an O^6^meG (Fig. **4c**) or AP site (Fig. **4d**) in double-stranded oligonucleotides and C opposite O^6^meG in a hairpin oligonucleotide (Fig. **4e**).

**Fig. 4:**
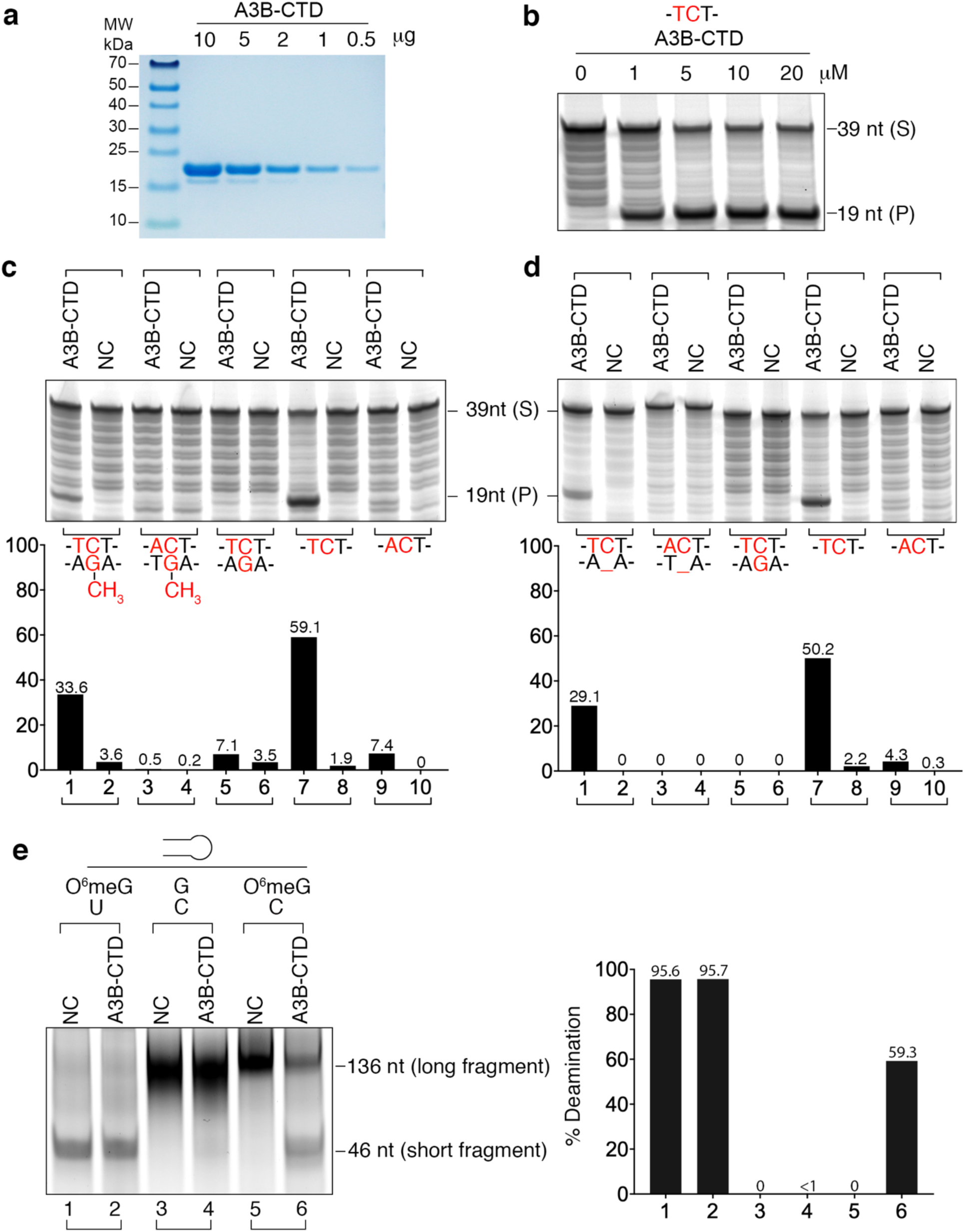
The A3B C-terminal domain (CTD) is sufficient to deaminate lesion-containing dsDNA in vitro. (**a**) Coomassie staining of purified A3B-CTD. (**b**) Deaminase activity of purified A3B-CTD (1, 5, 10, 20 µM) on TC-containing ssDNA. (**c,d**) *In vitro* deamination with A3B-CTD on O^6^meG/C (**c**) or AP/C (**d**) 39 bp oligonucleotides. Bar graphs (lower panels) show the amount of cleaved product relative to the amount of starting substrate. TC-containing ssDNA serves as a positive control, and AC-containing ssDNA and TC-containing dsDNA serve as negative controls. S, substrate; P, cleaved product. (**e**) *In vitro* deamination assay of A3B-CTD on O^6^meG/U, G/C, or O^6^meG/C-containing hairpin oligonucleotides. 20% urea polyacrylamide gel was stained with GelRed to visualize DNA. Bar graph (lower panel) shows the amount of cleaved product relative to the amount of starting substrate. NC, negative control, storage buffer used for A3B-CTD.

### A3B enhances MNNG-induced *γ*-H2AX foci in Hs578T cells by a deaminase independent mechanism

To determine the effect of A3B on O^6^meG/C-containing DNA in mammalian cells, we treated cells with an alkylating agent, MNNG, to generate O^6^meG lesions (Fig. **5a**). We used Hs578T breast cancer cells, because they contain low level of MGMT mRNA relative to three other breast cancer cell lines that we tested (Supplementary Fig. **S2**) and should be deficient in direct O^6^meG repair. Immuno-cytological assays (ICA) using an anti-O^6^meG antibody showed that O^6^meG content was increased in a dose-dependent manner in response to MNNG treatment (Fig. **5b,c**).

**Fig. 5:**
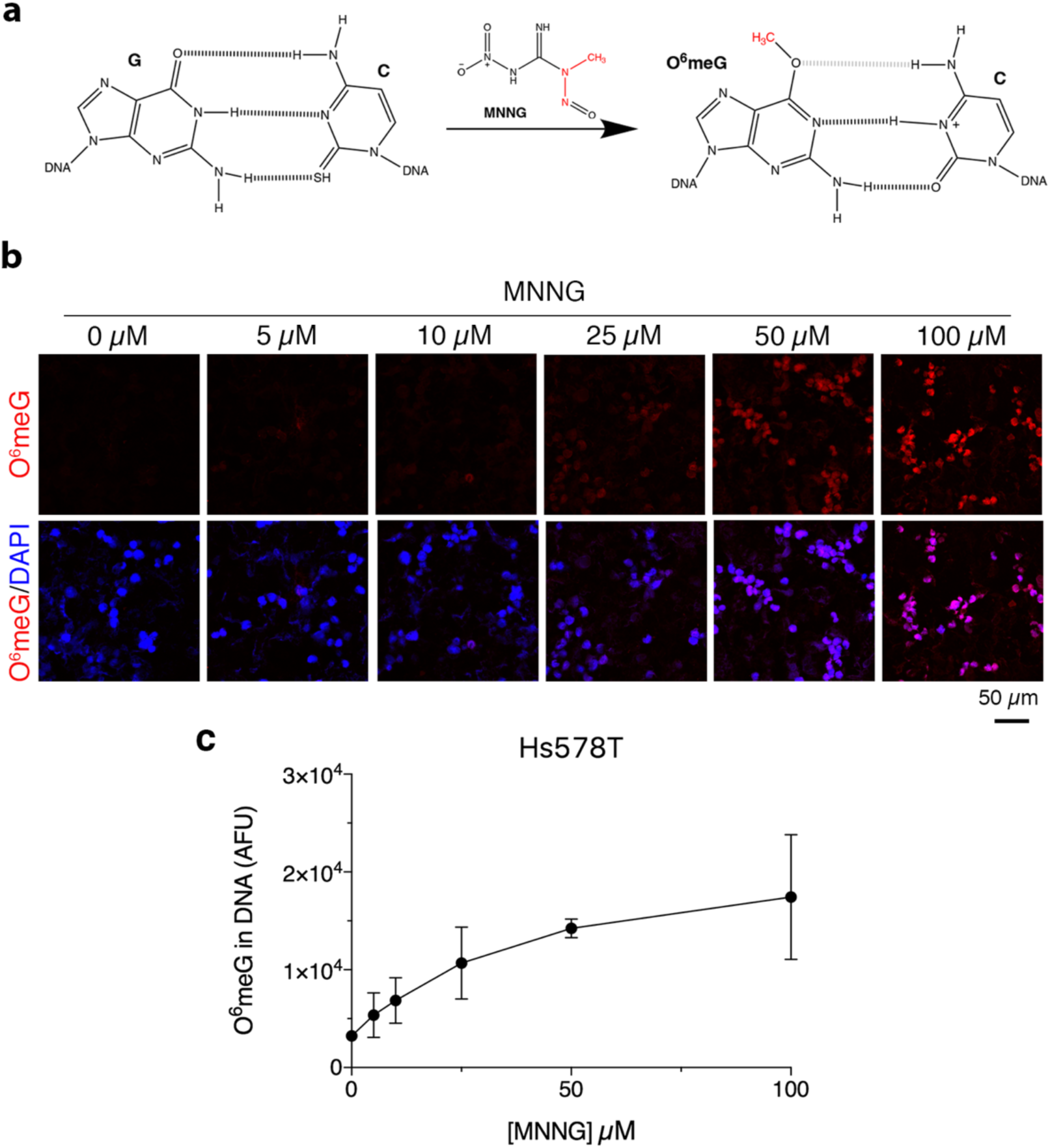
MNNG generates O^6^meG lesions in Hs578T cells. (**a**) Electrons on O^6^-G can attack the carbon of a methane diazonium cation (metabolic product of MNNG), resulting in the addition of an O^6^-methyl group on guanine. (**b**) Representative images of Hs578T cells stained with DAPI after an immunocytological assay (ICA) using EM 2-3 monoclonal antibody (targets O^6^meG residues) show that increasing concentration of MNNG increases the amount of O^6^meG. (**c**) Quantification of n = 6 fields per sample from 3 independent trials, error bars represent standard deviation.

MNNG can induce DSBs measured by γ-H2AX foci^21,36,37^. This occurs because replicative DNA polymerases can misread O^6^meG as an A and generate a T/O^6^meG mismatch, a substrate for mismatch repair (MMR)^21,38^. Persistence of O^6^meG in the single-stranded repair template generated during MMR initiates futile cycles of MMR (or BER) because O^6^meG is repeatedly copied into T, resulting in persistent single-stranded DNA, and eventual fork collapse and DSBs^21,31^. C paired to O^6^meG is susceptible to A3B deamination *in vitro* (Figs. **1d**, **2c**, **3**, **4c, 4e**), therefore we hypothesized that A3B may increase MNNG-induced damage by deaminating C to U, creating an MMR or BER substrate which would generate labile AP sites and eventual DSBs. A3B could also aggravate MNNG-induced DSBs by binding to exposed ssDNA^39^ and interfere with repair. We found that γ-H2AX lesions were increased in a dose-dependent manner in response to MNNG treatment (Supplementary Fig. **S3**).

To examine the effect of A3B on MNNG-induced γ-H2AX foci (Fig. **6a**), we transfected Hs578T cells with A3B-HA (Fig. **6b**, top) 48 h prior to MNNG treatment. Hs578T nuclear extracts contain substantial T<u>C</u>-specific deaminase activity (Fig. **6b**, lanes 1 and 2), which was increased approximately two-fold in cells that were transfected with the A3B-HA expression vector (Fig. **6b**, lanes 3 and 4). Immunofluorescence of γ-H2AX foci showed that A3B-HA expression significantly enhanced production of DSBs at sub saturating amounts of MNNG (<100 μM) in Hs578T cells (Fig. **6c,d** and Supplementary Fig. **S4a**) without affecting cell viability (Supplementary Fig. **S4b**).

**Fig. 6:**
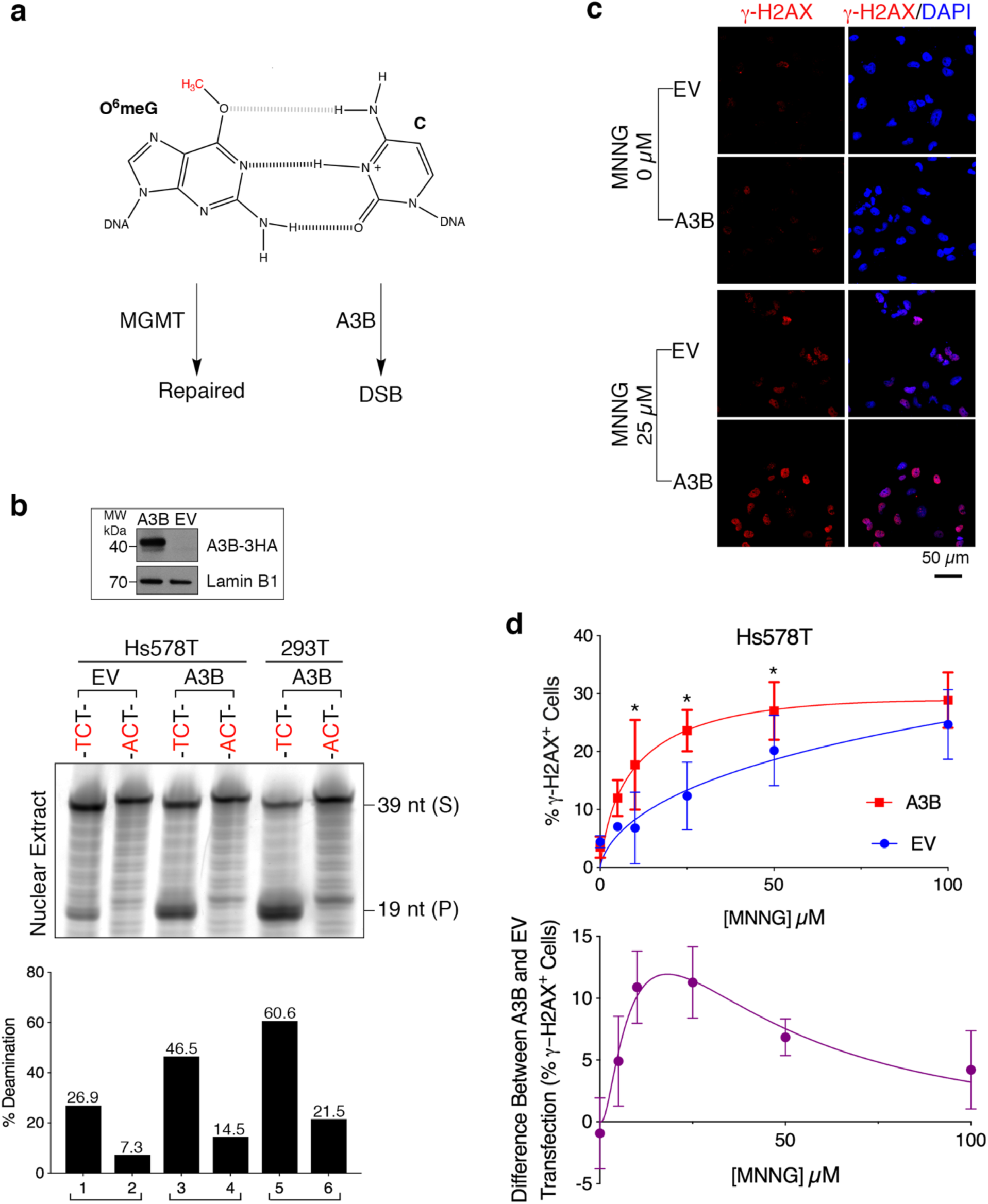
A3B enhances MNNG-induced γ-H2AX foci in Hs578T cells. (**a**) Proposed model of A3B-mediated MNNG-induced DSBs in cells. MGMT transfers the O^6^-methyl to its active site cysteine (Cys145), restoring the G/C base pair and inactivating MGMT. (**b**) Upper panel shows A3B-HA expression detected by western blot in Hs578T cells. LaminB1 serves as a loading control. *In vitro* deamination on 39 bp TC- or AC-containing ssDNA using Hs578T nuclear extracts transfected with an empty vector or A3B-HA expression vector (HEK293T extracts serve as a control). Bar graph shows the amount of cleaved product relative to the amount of starting substrate. (**c**) Representative images of Hs578T cells transfected with EV or A3B plasmids, treated with MNNG (25 µM), and stained with DAPI and Phospho-Histone H2A.X (Ser139) Antibody. The representative images for MNNG treatment at 5, 10, 50, and 100 µM are shown in Supplementary Fig. **S4**. EV, empty vector; A3B, A3B-3HA expression vector. (**d**) Percentage of cells with γ-H2AX positive foci (upper panel) in Fig. **6c** and Supplementary Fig. **S4a**. A3B enhances H2AX activation/phosphorylation in the presence, but not absence of MNNG. Bottom panel shows the difference between γ-H2AX positive foci percentages detected in cells transfected with A3B-HA and EV. n = 30 fields of view per sample from 3 independent trials, error bars represent standard deviation. Paired, two-tailed student’s t-tests confirm statistical significance, *p < 0.05 at 10, 25 and 50 µM MNNG treatments.

However, robust expression of two deaminase deficient A3B proteins (A3B-cat and T214D, Fig. **7a**, insert, lanes 5-8) also increased the production of γ-H2AX foci (generated at 25 μM MNNG) and to about the same extent as deaminase competent A3B-HA expression (Fig. **7b**), although the effect of A3B-cat was somewhat less. A3B-cat also contains a E68A mutation in the NTD, which may affect its binding to single strand DNA. Nonetheless, these results indicate that deaminase activity is dispensable for the effects mediated by A3B in MNNG-treated cells. Deaminase independent activity has been found in other instances of A3-mediated effects including A3B^40–45^. We also found that knockdown of endogenous A3B by siRNA did not affect the level of γ-H2AX foci (Supplementary Fig. **S5**). This result may not be surprising, given that endogenous levels of A3B in Hs578T cells is insufficient to generate observable γ-H2AX foci in the presence of MNNG at the higher fluorescence threshold we used in MNNG treated cells (see Materials and methods). At this threshold, even overexpression of A3B fails to produce a detectable increase of γ-H2AX foci (Fig. **6c,d**).

**Fig. 7:**
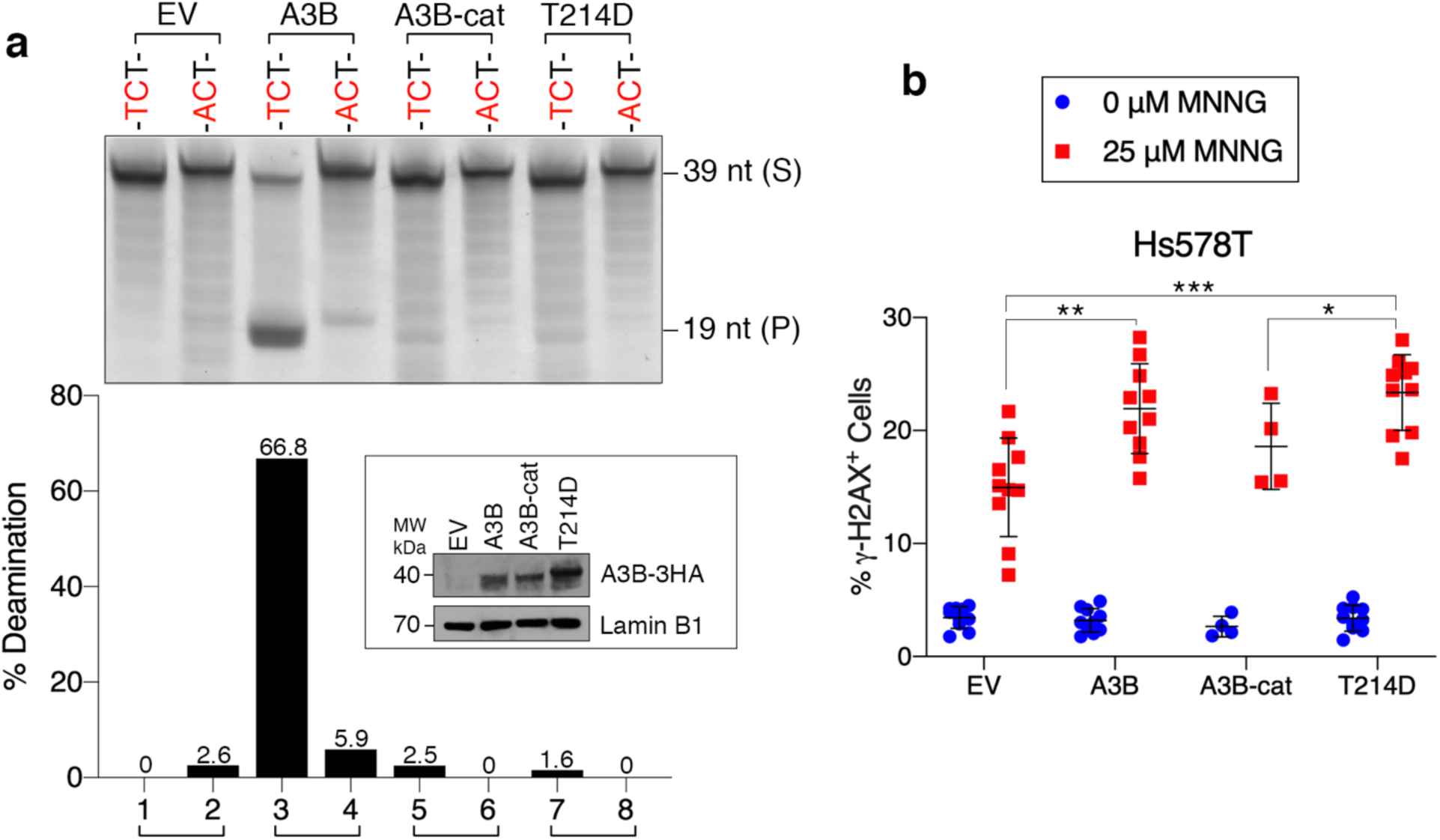
Effect of A3B deaminase activity on MNNG-induced γ-H2AX foci. (**a**) I*n vitro* deaminase activity of extracts of Hs578T cells transfected with the indicated expression vectors on TCT- or ACT-containing 39 nt ssDNA. Bar graph represents the amount of cleaved product produced by A3B-HA deamination relative to the amount of starting substrate. Insert shows the western blot of A3B-HA protein generated by the various expression vectors in Hs578T cells probed with anti-HA antibody. LaminB1 serves as a loading control. S, substrate; P, cleaved product. (**b**) γ-H2AX foci in Hs578T cells that had been transfected with wild-type or mutated A3B-HA expression vectors and treated with 25 µM MNNG (1 h) followed by a 1 h recovery. EV, empty vector; A3B-cat, double deaminase mutant (E68A, E255Q). n = 4 (A3B-cat), n = 10 (EV, A3B, T214D), line represents mean, error bars represent standard deviation. Two-tailed t-tests (equal variance) confirm statistical significance. *p < 0.05, **p < 0.01, ***p < 0.001.

### Fate of O^6^meG/C-containing plasmid DNA in Hs578T cells

To determine whether A3B expression affects mutations caused by O^6^meG/C in cells, we inserted an O^6^meG/C base pair (in a T<u>C</u>T or G<u>C</u>C context) into a shuttle vector^10^ (Supplementary Fig. **S6a**) and introduced it into Hs578T cells that had been transfected with either an A3B-HA expressing or empty vector (EV). After 24 h we isolated only those vectors that had replicated in Hs578T cells (see Materials and methods) and generated libraries from the 150 bp DNA flanking the inserted O^6^meG for paired-end DNA sequencing (Supplementary Table **S3**). If the O^6^meG-containing strand was replicated, O^6^meG would have been read as A most of the time, subsequently templating T. Therefore, we would have expected as much as half of the recovered plasmids to have an A at this site. However, the yield of A was less than 1.5% of the recovered products (Supplementary Fig. **6b** and Supplementary Table **S4**), indicating that most of the shuttle vector that we recovered resulted from replication of just the C-containing strand. Thus, most of the O^6^meG-containing strand likely suffered strand breakage due to futile rounds of MMR/BER and other processes, and was lost. As this kind of damage would produce DSBs on genomic DNA, the shuttle vector results are consistent with and corroborates the induction of genomic γ-H2AX foci by MNNG in Hs578T cells (Figs. **5** and **6**).

## Discussion

A3 deaminases target the N^4^ group of single-stranded C, with varying degrees of preference for the C in a T<u>C</u> motif, most highly by A3B. Here, we show that A3B can also deaminate C of T<u>C</u> opposite O^6^meG or AP sites in dsDNA (Figs. **1d**,**f**, **2c**, **3**, **4c-e**). Several proposed conformational structures and binding energy calculations show that the decreased O^6^-G/N^4^-C hydrogen bond strength weakens the O^6^meG/C base pair compared to a G/C base pair^20,46^. This weakened O^6^-G/N^4^-C hydrogen bond could expose the N^4^-C to A3 deaminases (Fig. **1a** and **6a**), although a significant amount of substrate may not be deaminated due to O^6^meG/C wobble base pairing between the N^1^-G and the N^4^-C hydrogen^46^. Likewise, deamination of C opposite an AP site may occur because the cytosine is free to rotate, thus providing an easier target for A3B.

Therefore, the weaker the hydrogen bonding involving the 4-amino group on C, the more likely it can be engaged by A3B. A crystal structure of the A3B-CTD and its substrate^32^ shows both the −1T (*i.e.*, T 5’ of C) and 0C are accommodated in the substrate binding pocket of the enzyme. As our results recapitulate the strong discrimination between the −1T vs the −1A (Fig. **1d,f** and **4c,d**), we assume that engagement of the 0C by the enzyme destabilizes the helix sufficiently to allow recognition of the −1 position, a result consistent with the known “breathing” dynamics of double-stranded nucleic acids^47–49^. At least 10,000 AP sites in addition to numerous O^6^meG adducts occur randomly throughout the genome and could well be a potential source of previously described random, isolated A3B mutations. Thus, our findings supplement observations of strand-coordinated clusters of A3B mutations observed in many human cancers and some non-cancerous tissues^6–8^.

The mode of action of MNNG is similar to other chemotherapeutic alkylating agents such as temozolomide and dacarbazine. They are metabolized to a methane diazonium cation, which can generate O^6^meG (Fig. **5a**) as well as other adducts^50^, leading to futile cycles of MMR and eventually DSBs. As expected, high levels of MGMT have been correlated with poor outcomes for such alkylating agent chemotherapy by repairing these genotoxic lesions^51–54^. As explained in the Results, we would expect that A3 deaminases could enhance the damage induced by MNNG, especially in the MGMT-deficient Hs578T cell line as the putative O^6^meG/C A3B substrate would not be directly repaired. A3B did act synergistically with MNNG in the production of DSBs (Fig. **6c,d**), but this effect was independent of its deaminase activity (Fig. **7b**).

A3B, like other members of the A3 family, is a single-strand specific DNA binding protein^44,55^. This property is presumably related to the numerous instances of A3 deaminase-independent effects^40–45^. Thus, our findings that A3B deaminase activity is not required for its aggravation of MNNG-induced DSBs suggest that it does so as a consequence of binding to the single-stranded regions that characterize MNNG-induced damage, perhaps by interfering with or delaying the eventual repair of these lesions.

We also examined the interaction between A3B and O^6^meG/C by inserting this base pair in either a T<u>C</u>T or G<u>C</u>C context on a plasmid shuttle vector, which replicates autonomously in breast cancer cells from its embedded SV40 origin (Supplementary Fig. **S6a**)^10^. Our results indicate that most of the O^6^meG-containing strand did not contribute to the replicated shuttle vector recovered in these experiments (Supplementary Fig. **S6b** and Supplementary Table **S4**), likely because it was irretrievably damaged during DNA repair by various means including futile cycles of MMR^21,38^.

That plasmid recovery was worse when O^6^meG is paired to C in a T<u>C</u>T context (*i.e.* an A3B context) than in a G<u>C</u>C context is consistent with the synergistic effect of A3B and MNNG treatment in the formation of DSBs (Fig. **6c,d**).

In summary, our biochemical evidence indicates that subtle but common genetic alterations of DNA structure can significantly affect protein-DNA interactions. Future studies could address the possibility that other APOBEC enzymes, or perhaps other proteins known to interact exclusively with ssDNA, behave in a similar manner to A3B when certain DNA lesions or higher order structures are introduced into double stranded DNA.

## Materials and methods

### Oligonucleotides

Oligonucleotides (excluding site-directed mutagenesis primers) were synthesized by Midland BioProducts or IDT and are listed in Supplementary Table **S1**. Oligonucleotides used for the *in vitro* deamination assay (39 nt) were annealed in equimolar ratios (100 mM Tris-HCl pH 7.5, 500 mM NaCl), heated at 95°C for 5 min and cooled to room temperature for 4 h. Annealing efficiency was monitored by native PAGE followed by Fujifilm FLA-5100 imaging on the 473 nm channel. AP sites were generated by treating oligonucleotides containing an internal uracil with UDG (NEB, M0280S) (0.08U/μL reaction) for 30 min at 37°C immediately before use. The 182 nt hairpin oligonucleotides were PAGE purified before use.

### Deamination assay

We performed deamination assays as described previously^35,56,57^. Immediately before performing the assay, cell or nuclear extracts were incubated with RNase A (1 μg/μL) for 15 min at 37°C. Then 25 μg (or 10 μM purified A3B-CTD) of the extract was incubated for 5 h at 37°C in a 40-80 μL reaction containing 500 nM oligonucleotide, 10 mM Tris-HCl pH 8.0, 50 mM NaCl, 1 mM DTT, 1 mM EDTA, 2 U UDG (NEB, M0280S), and 1x UDG buffer (NEB). We adjusted the reactions to 150 mM NaOH and incubated them for 20 min at 37°C, followed by 5 min at 95°C, then immediately chilled on ice, and in some cases, DNA was ethanol precipitated. Samples were heated for 5 min at 90°C in 1x NOVEX TBE-Urea Sample Buffer (Invitrogen), subjected to 12% or 20% 7.5 M Urea PAGE, and either imaged using the FUJIFILM FLA-5100 (FUJIFILM Life Science, 473 nm excitation) if fluorescein-labeled, or using GelRed (Biotium) in the case of hairpin oligonucleotides. Bands were quantified using Fiji software (NIH).

### Mammalian expression vectors

Mammalian expression vector pcDNA3.1(+) and phAPOBEC3B-HA were obtained respectively from Invitrogen and the NIH AIDS Reagent Program. phAPOBEC3B-HA has a pcDNA3 backbone with a 1275 bp KpnI/XhoI insert consisting of the hAPOBEC3B gene linked to 3 carboxy-terminal HA-tags.

### Cell lines and transfection

Breast cancer cell line Hs578T (HTB-126) was obtained from American Type Culture Collection (ATCC) and cultured as previously reported^58^. HEK293T cells (provided by Dr. Roland Owens, NIH) were maintained in DMEM with 10% FBS at 37°C in 5% CO_2_.

We transfected HEK293T cells with polyethylenimine (PEI, Polysciences)^59^ complexes of pcDNA3.1(+) or phAPOBEC3B-HA (6 µg per 10 cm dish) using the manufacturer’s protocol. We transfected Hs578T cells with pcDNA3.1(+), phAPOBEC3B-HA, or mismatch plasmids (1.5 µg per well in 6-well plate) using Lipofectamine 3000 Reagent (Thermo Fisher Scientific) following the manufacturer’s protocol.

### Preparation of mammalian cell extracts and western blotting

Incubation, extraction, and protein quantification were performed as previously described^35,56,60^. After plasmid transfection, cells were incubated for 48 h at 37°C in 5% CO_2_, harvested and lysed using M-PER Mammalian Protein Extraction Reagent (Thermo Scientific, 78501) (with 1x Roche Complete Protease Inhibitor and 100 mM NaCl). The supernatant was collected after a 10-min 16,200 *xg* centrifugation at 4°C and adjusted to 10% glycerol and centrifuged again for 10-min 16,200 *xg* at 4°C. We determined protein concentration with the PIERCE BCA Protein Assay Kit (Thermo Fisher Scientific).

Western blots were probed with anti-HA (1:1000, Rabbit polyclonal, Sigma-Aldrich, H6908) and anti-LaminB1 primary antibodies (1:2000, Rabbit polyclonal, abcam, ab16048) at 4°C overnight, followed by anti-Rabbit IgG-Peroxidase (1:10,000, Sigma-Aldrich, A0545) incubation for 1.5-2 h, and SUPERSIGNAL WEST PICO Chemiluminescent Substrate (Thermo Fisher Scientific) treatment for 5 min.

### Isolation of mammalian cell nuclei

We isolated nuclei as previously described^61^. Cells were resuspended in 5 volumes of extraction buffer A (20 mM Tris-HCl pH 7.5, 100 mM EDTA, 2 mM MgCl_2_), incubated for 2 min at room temperature, followed by 10 min on ice. We adjusted the extract to 1% NP-40, 1 mM PMSF, and 1x Complete Protease Inhibitor (Roche) and incubated the samples for 15 min on ice. We then passed the extracts through a 20-G needle and collected the nuclear pellet with a 10-min 500 *xg* centrifugation at 4°C. We carried out nuclear lysis, protein extraction, and quantification as described above.

### Construction of A3B deaminase mutants

We introduced mutations in phAPOBEC3B-HA using the Q5 Site-Directed Mutagenesis Kit (NEB) following the manufacturer’s protocol. Primers (synthesized by IDT) are listed in Supplementary Table **S2** and their annealing temperatures (T_a_) were calculated using NEBaseChanger. All constructs were confirmed by DNA sequencing (ACGT, Inc.). We also used the previously described double deaminase mutant A3B-cat (E68A, E255Q)^2,10^.

### A3B-CTD

Purified A3B-CTD protein (amino acid 187-378 of A3B, with mutations L230K and F308K, and an N-terminal SUMO tag cleaved off after purification) was a generous gift from Dr. Hideki Aihara’s laboratory at University of Minnesota and was expressed and purified as previously reported^32,33^. The protein was stored in 20 mM Tris-HCl (pH 7.4), 500 mM NaCl, and 5 mM *β*-mercaptoethanol.

### Chemicals

1-methyl-3-nitro-1-nitrosoguanidine (MNNG) was purchased from TCI America and dissolved in DMSO as a 1 M stock. 3-(4,5-dimethylthiazol-2-yl)-2,5-diphenyltetrazolium bromide (MTT) was purchased from Invitrogen and dissolved in 1x PBS (pH 7.4) as a 5 mg/mL stock.

### Immunocytological assay (ICA) to detect O^6^meG levels

An ICA was performed as previously described^62^. Hs578T cells were seeded in a 6-well plate, treated with MNNG as described above, and dotted on Superfrost Plus Gold Slides (Thermo Fisher Scientific). Cells were fixed at −20°C in cooled methanol for 30 min, washed with 1x PBS, treated with an alkali/methanol solution (60% 70 mM NaOH/140 mM NaCl, 40% methanol) for 5 min on ice to lyse the cell membranes, and washed with 1x PBS. Slides were placed in a moist chamber for the remainder of the protocol. They were treated with 540 µg/mL Pepsin (Sigma, P6887) (activated with 20 mM HCl immediately prior to use) for 10 min at 37°C. Slides were washed with 1x PBS and treated with 800 µg/mL Proteinase K (Thermo Fisher Scientific, EO0491) for 10 min at 37°C, washed in PBS-glycine (0.2% glycine) for 10 min, blocked in 5% skim milk (in 1x PBS) for 30 min, and stained with 100 µg/mL EM 2-3 monoclonal antibody (Squarix, SQM003.1) (1:300 in 5% BSA) at 4°C overnight. Following washes with PBS-T (0.25% Tween 20) and 1x PBS, cells were stained with Alexa Fluor 568 Goat anti-Mouse IgG (H+L) (Thermo Fisher Scientific, A-11004) (1:400 in 5% BSA) in a dark moist chamber for 75 min. The cells were washed as above and stained with 1 µg/mL DAPI for 30 min and washed in 1x PBS. IMMUNOSELECT Antifading Mounting Medium (Squarix) was applied and coverslips were sealed with nail polish and incubated at 4°C overnight. Slides were imaged with a KEYENCE Digital Microscope using 1/28 and 1/30 second exposure time for blue and red channels, respectively, and 2 fields per sample were analyzed using Fiji software. All cells and 10 background readings in each field of view were measured for area, integrated density, and mean gray values. To calculate corrected total cell fluorescence (CTCF) of each cell, the cell area was multiplied by the average mean gray value of the 10 background readings, and the resulting value was subtracted from the cell raw integrated density. The average CTCF was calculated by averaging CTCF values for all cells from both fields.

### γ-H2AX staining

Hs578T cells were transfected with pcDNA3.1(+) or phAPOBEC3B-HA, and after 32 h transferred to a NUNC LABTEK II Chamber Slide System (Thermo Fisher Scientific). After 16 h, the cells were incubated with MNNG for 1 h at 37°C in complete media, washed twice with 1x PBS, and incubated in fresh media for 1 h at 37°C without MNNG (recovery phase). Cells were washed twice in 1x PBS, fixed with 4% formaldehyde (ThermoFisher Scientific, diluted in 1x PBS) for 15 min at 37°C, followed by permeabilization with 0.1% Triton xX-100 (in 1x PBS) for 10 min at room temperature. After washing twice in 1x PBS, cells were blocked with 1% BSA (in 1x PBS) for 30 min at room temperature and incubated overnight at 4°C with Phospho-Histone H2A.X (Ser139) Antibody (Cell Signaling, 2577) in 1% BSA. Cells were blocked again and incubated for 75 min at room temperature with Goat anti-Rabbit IgG (H+L) Cross-Absorbed Secondary Antibody, Alexa Fluor 568 (Thermo Fisher Scientific) (in 1% BSA). The chambers were disassembled, and slides with cells were mounted with PROLONG DIAMOND Antifade Mountant with DAPI (Thermo Fisher Scientific). After incubating overnight, slides were imaged with a Keyence Digital Microscope using 1/15 and 1/12 second exposure time for blue and red channels, respectively. Analysis was performed using Fiji software on 10 random fields per sample. Color thresholding for the red channel was set to determine γ-H2AX^+^ cells (moments, brightness 50, convert image to binary, count cells ≥ 150 pixels). The number of red-stained nuclei was divided by the number of DAPI-stained nuclei and multiplied by 100 to obtain the percentage of γ-H2AX positive cells.

### siRNA knockdown of A3B

We knocked down endogenous A3B in Hs578T cells using 10 nM siRNA (Dharmacon, J-017322-08-0005) and Lipofectamine RNAiMAX Transfection Reagent (Thermo Fisher Scientific) following the manufacturer’s protocol. We used scrambled siRNA (Dharmacon, UAGCGACUAAACACAUCAA, siScr) as a negative control.

### qRT-PCR

We extracted mRNA from breast cancer cells using the PURELINK RNA Mini Kit (Life Technologies) and synthesized cDNA with the SUPERSCRIPT III First-Strand Synthesis System (Thermo Fisher Scientific). We carried out qRT-PCR with TaqMan Universal PCR Master Mix, no AmpErase UNG (Thermo Fisher Scientific) on a STEPONEPLUS Real-Time PCR System (Applied Biosystems). The original data for mRNA level of MGMT and housekeeping genes in four breast cancer cells shown in Supplementary Fig. **S2** was adapted from Shen et al^58^.

### MTT assay

We performed an MTT assay^63^ to determine cell viability of MNNG treated Hs578T cells. Metabolically active cells can reduce MTT (yellow) to formazan (purple). 48 h after Hs578T cells were transfected with pcDNA3.1(+) or phAPOBEC3B-HA, they were incubated in a 96-well plate with MNNG for 16 h at 37°C in 5% CO_2_, washed with 1x PBS, and treated with 500 μg/mL MTT for 4 h at 37°C in 5% CO_2_. Media was removed, DMSO was added to resuspend cells, and absorbance was measured at 595 nm using the Bio-Rad Model 680 Microplate Reader.

### Construction of O^6^meG/C-containing plasmids and processing in Hs578T cells

O^6^meG/C base pairs were introduced into an SV40-based shuttle vector^64^ in both a T<u>C</u>T and G<u>C</u>C context as previously described^10^. Briefly, 120U Nt.BbvCI (NEB) was used to nick 60 µg of plasmid in the mismatch region of pSP189-FM1 at 2 positions 39 bp apart on the same strand at 37°C overnight. The nicked strand was removed by annealing to a complementary biotinylated oligonucleotide (Supplementary Table **S1**) for 1 h at 37°C, and the resulting duplex was captured on 3 mg streptavidin-coated magnetic beads (Roche, rotation for 2 h at 37°C). After extraction with phenol-chloroform and precipitation with ethanol, 4 μg of the gapped plasmid was annealed to the appropriate 39 nt oligonucleotides (1:100 molar ratio, Supplementary Table **S1**) and ligated overnight with T4 DNA Ligase (NEB) at 16°C. Restoration of a KpnI restriction site present in the gapped region confirmed incorporation of mismatch oligonucleotides. Immediately before transfection into Hs578T cells, 4 µg of plasmid was treated with 1U Klenow Fragment (3’ to 5’ exo-, NEB) for 10 min at 37°C to fill any remaining gapped plasmid^10^. The O^6^meG/C-containing plasmids were transfected into Hs578T cells that had been transfected with an A3B expression vector (phAPOBEC3B-HA) 40 h earlier. After 24 h, cells were harvested, and plasmids were extracted using the WIZARD PLUS SV Minipreps DNA Purification System (Promega). Plasmids were then treated with 5U DpnI (NEB) for 30 min at 37°C to degrade any non-replicated plasmids, and used as template DNA for NGS sample preparation using Q5 High-Fidelity DNA Polymerase (NEB) (primers shown in Supplementary Table **S3**). The resulting 287 bp PCR products (amplicons) containing Illumina adapters and the 151 bp of plasmid DNA that flanked the mismatch region were purified using the QIAQUICK PCR Purification Kit (Qiagen) and quantified with PicoGreen (Thermo Fisher Scientific). Equimolar amounts of pooled amplicons were subjected to paired end 2 x 150 deep sequencing (ACGT, Inc.).

## Acknowledgements

We thank Dr. Judith Levin and Dr. Tiyun Wu (Section on Viral Gene Regulation, NICHD, NIH) for assistance in optimizing the *in vitro* deamination assay, Dr. Charles Jones (LCMB, NIDDK, NIH) for assisting with site-directed mutagenesis, and Dr. Jürgen Thomale (Center for Medical Biotechnology, University of Essen) for generous technical support with the ICA. We greatly thank Dr. Hideki Aihara (College of Biological Sciences, University of Minnesota) for providing the purified A3B-CTD protein. The A3B-cat expression vector was constructed in this lab by Dr. Jia Chen^12^ (present address, School of Life Science and Technology, ShanghaiTech University).

## Author contributions

J.H.C. proposed the idea of C paired to O^6^meG as a possible A3B substrate, and the rest of the authors participated in experimental design, data acquisition and interpretation. J.H.C., M.F.C., and A.V.F. wrote the manuscript, G.M.T. and B.S. helped in the editing and generation of figures.

