## Supplementary Information for "The effect of APOBEC3B deaminase on double-stranded DNA"

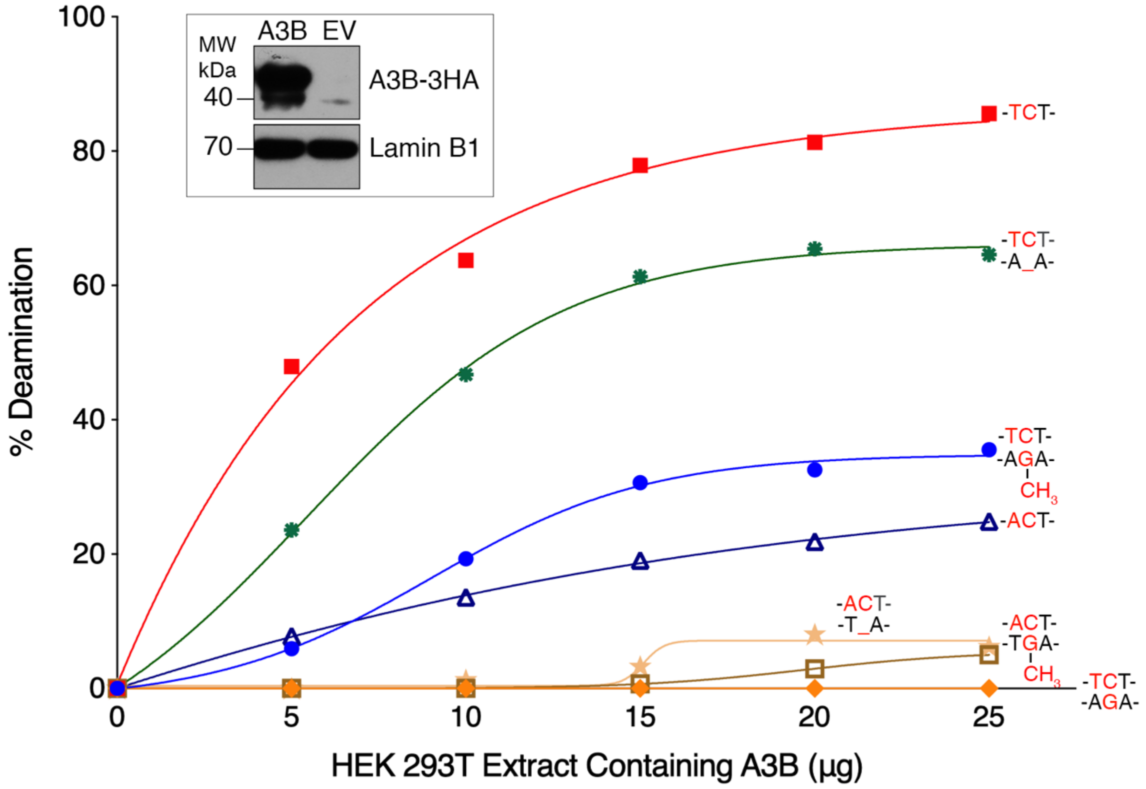


**Supplementary Fig. S1**: *Deaminase activity of HEK293T whole cell extract from cells transfected with A3B-HA expression vector.* Western blot shows A3B-HA expression using anti-HA antibody. LaminB1 serves as a loading control (insert). Curves represent the deaminase activity on the indicated 39 bp oligonucleotide substrates.

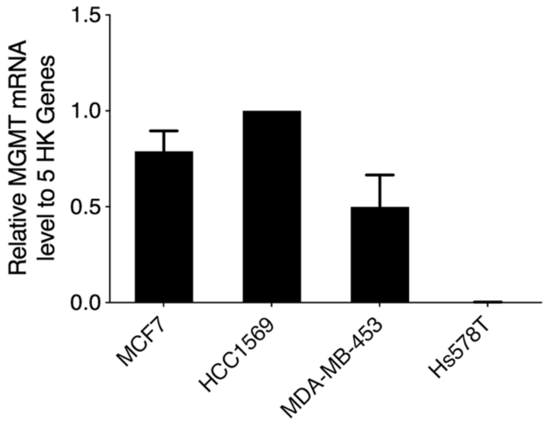


**Supplementary Fig. S2**: *Hs578T cell line is deficient in MGMT mRNA*. qRT-PCR determined MGMT mRNA levels in 4 breast cancer cell lines normalized to 5 housekeeping genes (ACTB, B2M, GAPDH, HPRT1, RPLP0) (The origin of the data is described in Materials and Methods/qRT-PCR). $n \geq 3$, error bars represent standard deviation. A two-sample equal variance, two-tailed t-test confirms statistical significance.

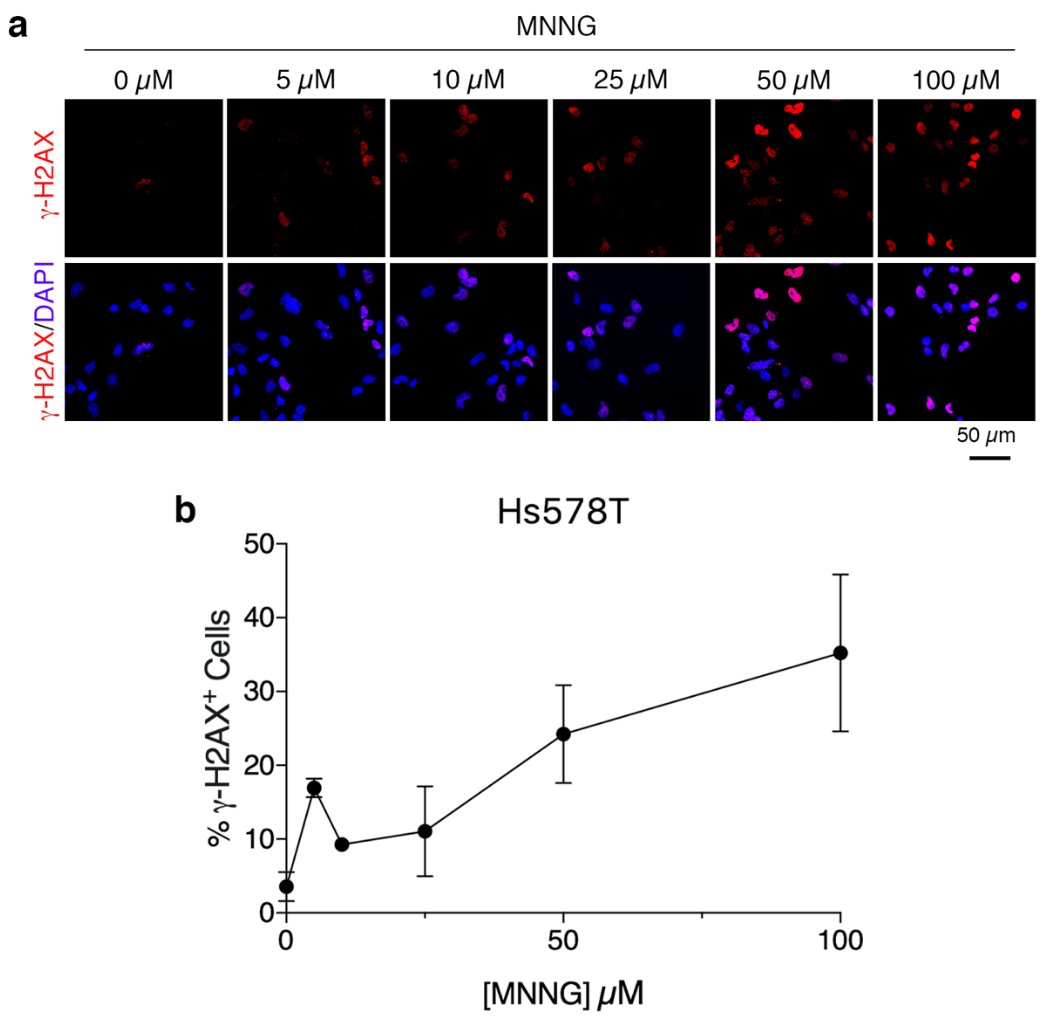


**Supplementary Fig. S3**: *MNNG induces* γ-*H2AX foci in Hs578T cells.* (**a**) Representative images of Hs578T cells stained with DAPI and Phospho-Histone H2A.X (Ser139) Antibody after MNNG treatment (1 h) followed by a 1 h recovery. Increasing MNNG concentration increases amount of γ-H2AX foci. (**b**) Quantification of (**a**) using n = 20 fields per sample from 2 independent trials, error bars represent standard deviation.

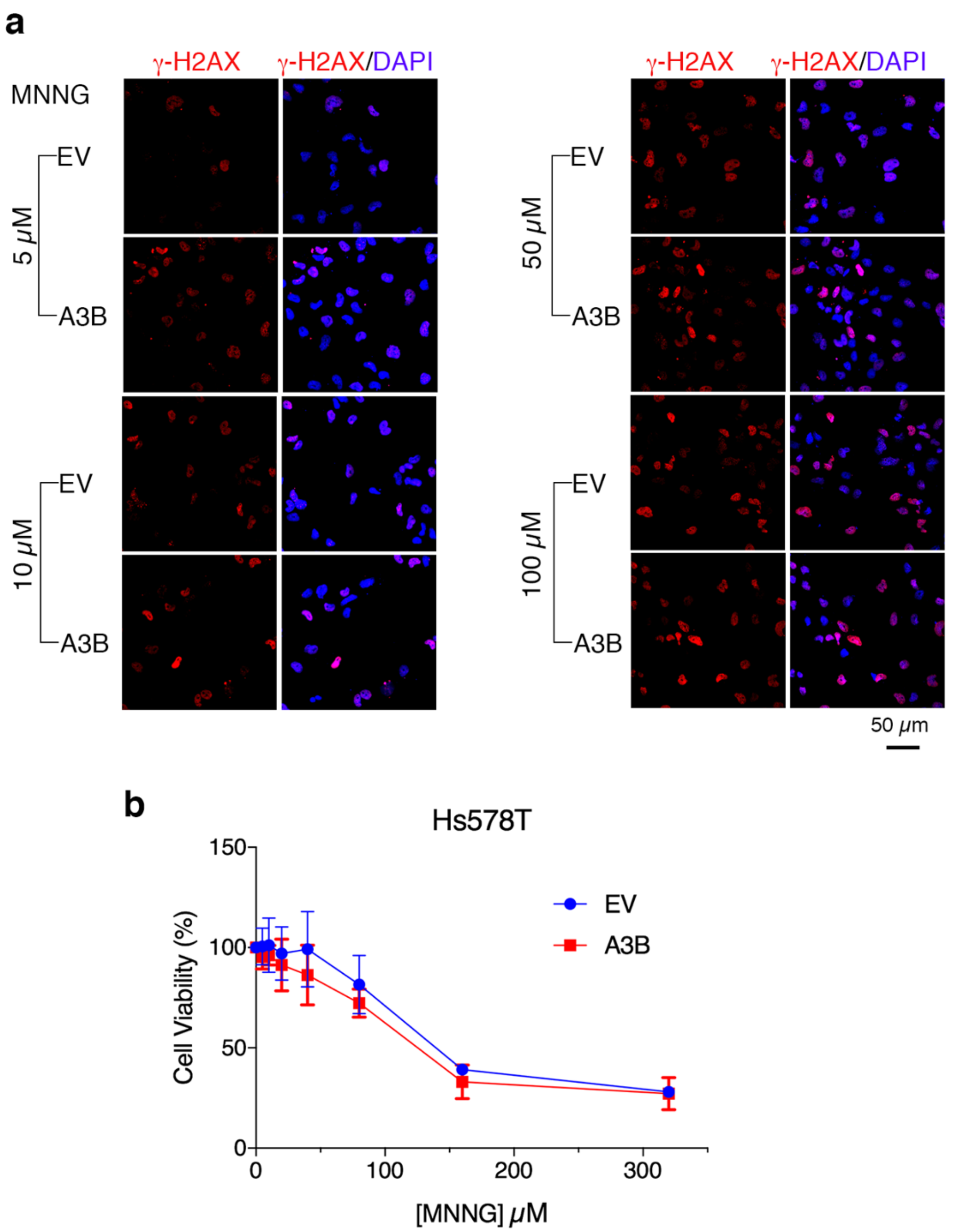


**Supplementary Fig. S4**: *A3B enhances MNNG-induced γ-H2AX foci in Hs578T cells without affecting cell viability*. (**a**) Representative images of Hs578T cells stained with DAPI and Phospho-Histone H2A.X (Ser139) Antibody show that A3B enhances H2AX activation/phosphorylation in the presence but not absence of MNNG. A3B, A3B-HA expression vector; EV, empty vector. 0 μM and 25 μM MNNG-treated cells are shown in Fig. **6c**. (**b**) MTT assay shows decreased cell metabolic activity with increasing MNNG concentration (16 h treatment), but no significant change when A3B-HA is introduced. n = 9 from 3 independent trials, error bars represent standard deviation.

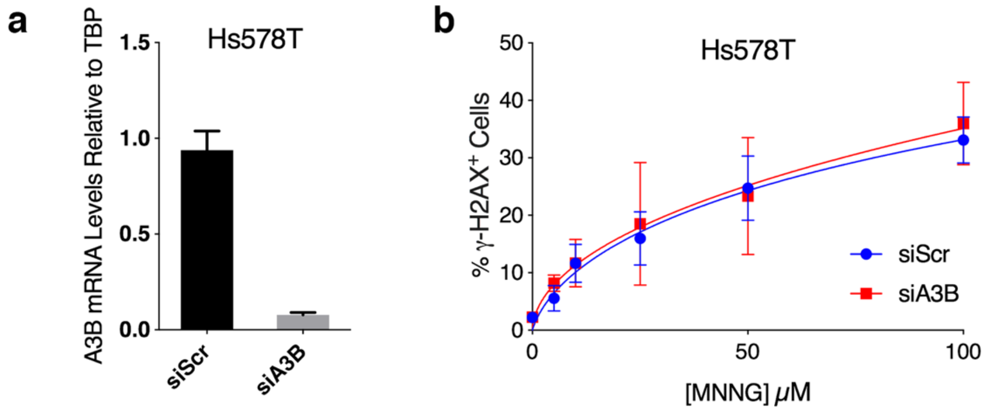


**Supplementary Fig. S5**: *Knockdown of A3B transcripts does not affect MNNG-induced γ-H2AX foci in Hs578T cells.* (**a**) qRT-PCR shows A3B mRNA levels are decreased in Hs578T cells transfected with siA3B compared to scrambled siRNA (siScr). n = 3 technical replicates from 1 trial, error bars represent standard deviation. (**b**) Hs578T cells treated with MNNG show that knockdown of endogenous A3B does not affect γ-H2AX foci. n = 30 fields per sample from 3 independent trials, error bars represent standard deviation.

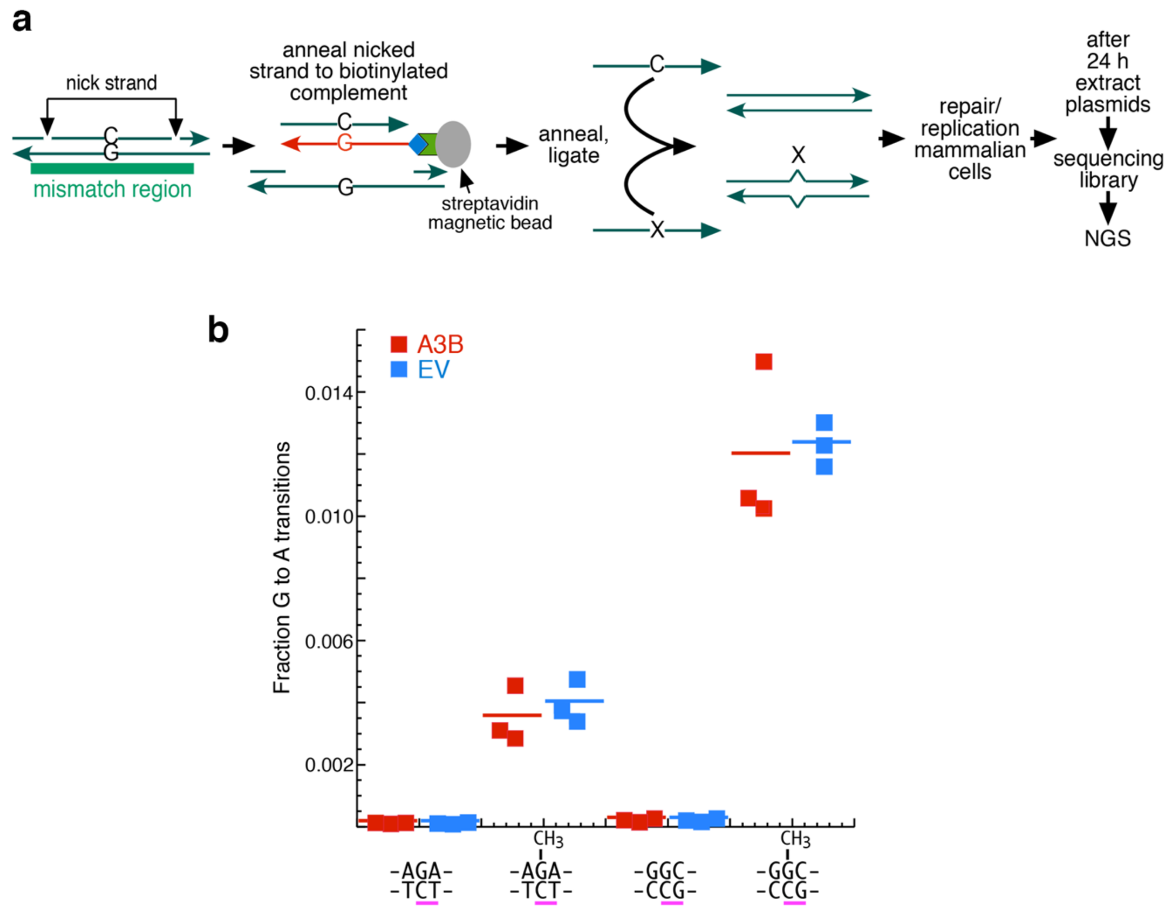


**Supplementary Fig. S6**: *Fate of $O^6$meG/C base pair in a shuttle vector in Hs578T cells.* (**a**) Schematic of mismatch plasmid construction. See MATERIALS AND METHODS for details. (**b**) Recovery of G to A transitions of $O^6$meG paired to C in a TCT context occurs one third as often compared to when the C is in a GCC context. In all conditions, transversions (G-to-T and G-to-C) consisted of <0.1% of mutations. Therefore, only G-to-A transitions were plotted. (Supplementary Table **S4**). Horizontal line represents mean. A paired, two-tailed student's t-test confirms statistical significance.

**Supplementary Table S1:** Oligonucleotides used for *in vitro* deamination assay and $O^6$meG/C plasmid construction

| Name | Sequence |
|---|---|
| O6G/TC | 5'-ATTATTATTATTATTATTA{O6-Methyl-dG}AATTATTATTATTATTATT-3'<br>3'-TAATAATAATAATAATAAT**C**-------------**T**TAATAATAATAATAATAA-{f}-5' |
| O6G/AC | 5'-ATTATTATTATTATTATTA{O6-Methyl-dG}TATTATTATTATTATTATT-3'<br>3'-TAATAATAATAATAATAAT**C**-------------**A**TAATAATAATAATAATAA-{f}-5' |
| G/TC | 5'-ATTATTATTATTATTATTAGAATTATTATTATTATTATT-3'<br>3'-TAATAATAATAATAATAAT**CT**TAATAATAATAATAATAA-{f}-5' |
| TC | 5'-{f}-AATAATAATAATAATAAT**TC**TAATAATAATAATAATAAT-3' |
| AC | 5'-{f}-AATAATAATAATAATAAT**AC**TAATAATAATAATAATAAT-3' |
| AP/TC | 5'-ATTATTATTATTATTATTA{dU}AATTATTATTATTATTATT-3'<br>3'-TAATAATAATAATAATAAT**C**---**T**TAATAATAATAATAATAA-{f}-5' |
| AP/AC | 5'-ATTATTATTATTATTATTA{dU}TATTATTATTATTATTATT-3'<br>3'-TAATAATAATAATAATAAT**C**---**A**TAATAATAATAATAATAA-{f}-5' |
| G/TC-1 | 5'-CTGCTACTGCTACTGCTACGAAATAATAATAATAATAAT-3'<br>3'-GACGATGACGATGACGATG**CT**TTATTATTATTATTATTA-{f}-5' |
| AP/TC-1 | 5'-CTGCTACTGCTACTGCTA{dU}GAAATAATAATAATAATAAT-3'<br>3'-GACGATGACGATGACGATG---**CT**TTATTATTATTATTATTA-{f}-5' |
| TC-1 | 5'-{f}-ATTATTATTATTATTATT**TC**GTAGCAGTAGCAGTAGCAG-3' |
| G/C | 5'-TCAGCTGACGTCGGACGGAGGCGGCAGAACCTAGGTACC-3' |
| O6G/TC | 5'-TCAGCTGACGTCGGACGGAGGCGGCA{O6-Methyl-G}AACCTAGGTACC-3' |
| O6G/GC | 5'-TCAGCTGACGTCGGACGGAGGCG{O6-Methyl-G}CAGAACCTAGGTACC-3' |

{f}, fluorescein; ---, inserted for alignment; TC and AC underlined in bold

**Supplementary Table S2:** Primers used for site-directed mutagenesis

| Name | Forward Primer | Reverse Primer | $T_a$ (°C) |
|---|---|---|---|
| T214D | ACGGCGCCAG*gac*TACTTGTGCT | CGAAGGACCAAAGGGTCATTATTAAA | 66 |
| H253S | TTACGGCCGC*agt*GCGGAGCTGC | AAGCCACAGAGAAGATTCTTAGC | 65 |
| A254P | CGGCCGCCAT*ccc*GAGCTGCGCT | TAAAAGCCACAGAGAAGATTCTTAGCCTC | 69 |
| A254G | CGGCCGCCAT*ggt*GAGCTGCGCT | TAAAAGCCACAGAGAAGATTCTTAGCC | 68 |
| S282A | CATCTCCTGG*gcc*CCCTGCTTCTC | AACCAAGTGACCCTGTAG | 62 |
| C284G | CTGGAGCCCC*ggt*TTCTCCTGGG | GAGATGAACCAAGTGACCCTG | 66 |
| C289G | CTCCTGGGGC*gga*GCCGGGGAAG | AAGCAGGGGCTCCAGGAG | 71 |
| Y313A | TGCCCGCATC*gcc*GATTACGACCCC | GCGAAGATGCGCAGTCTC | 64 |

Lower case letters code for the amino acid substitution

**Supplementary Table S3:** Primer indices used for shuttle vector NGS

| Name | i5 index | i7 index |
|---|---|---|
| EV_G/C_1 | TATAGCCT | ATCGTGAT |
| EV_G/C_2 | TATAGCCT | ATACATCG |
| EV_G/C_3 | TATAGCCT | ATGCCTAA |
| A3B_G/C_1 | GGCTCTGA | ATCGTGAT |
| A3B_G/C_2 | GGCTCTGA | ATACATCG |
| A3B_G/C_3 | GGCTCTGA | ATGCCTAA |
| EV_O6G/TC_1 | ATAGAGGC | ATCGTGAT |
| EV_O6G/TC_2 | ATAGAGGC | ATACATCG |
| EV_O6G/TC_3 | ATAGAGGC | ATGCCTAA |
| A3B_O6G/TC_1 | AGGCGAAG | ATCGTGAT |
| A3B_O6G/TC_2 | AGGCGAAG | ATACATCG |
| A3B_O6G/TC_3 | AGGCGAAG | ATGCCTAA |
| EV_O6G/GC_1 | CCTATCCT | ATCGTGAT |
| EV_O6G/GC_2 | CCTATCCT | ATACATCG |
| EV_O6G/GC_3 | CCTATCCT | ATGCCTAA |
| A3B_O6G/GC_1 | TAATCTTA | ATCGTGAT |
| A3B_O6G/GC_2 | TAATCTTA | ATACATCG |
| A3B_O6G/GC_3 | TAATCTTA | ATGCCTAA |

Full primer sequences

F_primer:AATGATACGGCGACCACCGAGATCTACAC[i5]ACACTCTTTCCCTACACGACGCTCTTCCGATCTGAATCCTTCCCCTCTAAC

R_primer:CAAGCAGAAGACGGCATACGAGAT[i7]GTGACTGGAGTTCAGACGTGTGCTCTTCCGATCTACGCGGCCCGACCTCGAC

**Supplementary Table S4:** Fate of $O^6$meG/C after replication on a shuttle vector in Hs578T cells

| NAME | P | R | TRI | GRF | A | C | G | T | TO | TR | TV | NUM | TO/NUM | A/NUM | E |
|---|---|---|---|---|---|---|---|---|---|---|---|---|---|---|---|
| **A3B_G/TC_1** | 76 | G | AGA | 64 | 42 | 9 | 0 | 13 | 64 | 42 | 22 | 309590 | 0.0002 | | |
| **A3B_G/TC_2** | 76 | G | AGA | 51 | 34 | 4 | 0 | 13 | 51 | 34 | 17 | 298217 | 0.0002 | | |
| **A3B_G/TC_3** | 76 | G | AGA | 51 | 36 | 7 | 0 | 8 | 51 | 36 | 15 | 258976 | 0.0002 | | |
| **A3B_G/GC_1** | 73 | G | GGC | 108 | 55 | 11 | 0 | 42 | 108 | 55 | 53 | 309590 | 0.0003 | | |
| **A3B_G/GC_2** | 73 | G | GGC | 72 | 30 | 11 | 0 | 31 | 72 | 30 | 42 | 298217 | 0.0002 | | |
| **A3B_G/GC_3** | 73 | G | GGC | 78 | 39 | 5 | 0 | 34 | 78 | 39 | 39 | 258976 | 0.0003 | | |
| **A3B_O6G/TC_1** | 76 | G | AGA | 1578 | 1560 | 3 | 0 | 15 | 1578 | 1560 | 18 | 340507 | 0.0046 | 0.0046 | 0.5 |
| **A3B_O6G/TC_2** | 76 | G | AGA | 1009 | 985 | 4 | 0 | 20 | 1009 | 985 | 24 | 343340 | 0.0029 | 0.0029 | 0.5 |
| **A3B_O6G/TC_3** | 76 | G | AGA | 969 | 948 | 6 | 0 | 15 | 969 | 948 | 21 | 304212 | 0.0032 | 0.0031 | 0.5 |
| **A3B_O6G/GC_1** | 73 | G | GGC | 5473 | 5410 | 9 | 0 | 54 | 5473 | 5410 | 63 | 363387 | 0.0151 | 0.0149 | 0.5 |
| **A3B_O6G/GC_2** | 73 | G | GGC | 3448 | 3406 | 13 | 0 | 29 | 3448 | 3406 | 42 | 333524 | 0.0103 | 0.0102 | 0.5 |
| **A3B_O6G/GC_3** | 73 | G | GGC | 3269 | 3227 | 8 | 0 | 34 | 3269 | 3227 | 42 | 306465 | 0.0107 | 0.0105 | 0.5 |
| **EV_ G/TC_1** | 76 | G | AGA | 80 | 52 | 11 | 0 | 17 | 80 | 52 | 28 | 349515 | 0.0002 | | |
| **EV_ G/TC_2** | 76 | G | AGA | 63 | 40 | 5 | 0 | 18 | 63 | 40 | 23 | 327444 | 0.0002 | | |
| **EV_ G/TC_3** | 76 | G | AGA | 49 | 28 | 1 | 0 | 20 | 49 | 28 | 21 | 295435 | 0.0002 | | |
| **EV_ G/GC_1** | 73 | G | GGC | 100 | 45 | 14 | 0 | 41 | 100 | 45 | 55 | 349515 | 0.0003 | | |
| **EV_ G/GC_2** | 73 | G | GGC | 82 | 37 | 10 | 0 | 35 | 82 | 37 | 45 | 327444 | 0.0003 | | |
| **EV_ G/GC_3** | 73 | G | GGC | 103 | 61 | 11 | 0 | 31 | 103 | 61 | 42 | 295435 | 0.0003 | | |
| **EV_ O6G/TC_1** | 76 | G | AGA | 1707 | 1682 | 6 | 0 | 19 | 1707 | 1682 | 25 | 353355 | 0.0048 | 0.0048 | 0.5 |
| **EV_ O6G/TC_2** | 76 | G | AGA | 1180 | 1157 | 5 | 0 | 18 | 1180 | 1157 | 23 | 308866 | 0.0038 | 0.0037 | 0.5 |
| **EV_ O6G/TC_3** | 76 | G | AGA | 835 | 815 | 0 | 0 | 20 | 835 | 815 | 20 | 239781 | 0.0035 | 0.0034 | 0.5 |
| **EV_ O6G/GC_1** | 73 | G | GGC | 4399 | 4346 | 8 | 0 | 45 | 4399 | 4346 | 53 | 335898 | 0.0131 | 0.0129 | 0.5 |
| **EV_ O6G/GC_2** | 73 | G | GGC | 3658 | 3613 | 10 | 0 | 35 | 3658 | 3613 | 45 | 313218 | 0.0117 | 0.0115 | 0.5 |
| **EV_ O6G/GC_3** | 73 | G | GGC | 3300 | 3250 | 9 | 0 | 41 | 3300 | 3250 | 50 | 266910 | 0.0124 | 0.0122 | 0.5 |

Only the rows corresponding to the location of $O^6$meG are shown: P, position of G in the reference (R) sequence; TRI, its trinucleotide context; GRF, the number of times the reference G was mutated; A, C, G, T, the mutational fate of GRF; TO, TR, TV, total mutations, transitions, transversions; NUM, number of paired sequences analyzed; E, the expected fraction of A's recovered if all of the $O^6$MeG's were replicated or repaired (see text). A/NUM are the values plotted in Supplementary Fig. **6**.
